# Expression patterns of *Plasmodium falcip*arum clonally variant genes at the onset of a blood infection in non-immune humans

**DOI:** 10.1101/2021.02.23.432621

**Authors:** Anastasia K. Pickford, Lucas Michel-Todó, Florian Dupuy, Alfredo Mayor, Pedro L. Alonso, Catherine Lavazec, Alfred Cortés

## Abstract

Clonally variant genes (CVGs) play fundamental roles in the adaptation of *Plasmodium falciparum* parasites to the fluctuating conditions of the human host. However, their expression patterns under the natural conditions of the blood circulation have been characterized in detail only for a few specific gene families. Here we provide a detailed characterization of the complete *P. falciparum* transcriptome across the full intraerythrocytic development cycle (IDC) at the onset of a blood infection in non-immune human volunteers. We found that the vast majority of transcriptional differences between parasites obtained from the volunteers and the parental parasite line maintained in culture occur in CVGs. Specifically, we observed a major increase in the transcript levels of most members of the *pfmc-2tm* and *gbp* families and of specific genes of other families, in addition to previously reported changes in *var* and *clag3* genes expression. Large transcriptional differences correlate with changes in the distribution of heterochromatin, confirming their epigenetic nature. The analysis of parasites collected at different time points along the infection indicates that when parasites pass through transmission stages, the epigenetic memory at CVG loci is lost, resulting in a reset of their expression state and reestablishment of new epigenetic patterns.

**Importance:** The ability of malaria parasites to adapt to changes in the human blood environment, where they produce long term infection associated with clinical symptoms, is fundamental for their survival. Clonally variant genes, regulated at the epigenetic level, play a major role in this adaptive process, as changes in the expression of these genes result in antigenic and functional alterations that enable immune evasion and provide phenotypic plasticity. However, the way these genes are expressed under the natural conditions of the human circulation or how their expression is affected by passage through transmission stages is not well understood. Here we provide a comprehensive characterization of the expression patterns of these genes at the onset of human blood infections, which reveals major differences with *in vitro* cultured parasites and also distinctive alterations between different families of clonally variant genes. We also show that epigenetic patterns are erased and reestablished during transmission stages.

## Introduction

*Plasmodium spp.* are responsible for the globally important disease malaria, which causes over 200 million cases and almost half a million deaths per year (1). Their complex life cycle is split between different hosts and cell types. The invertebrate mosquito host injects the infective parasite forms known as sporozoites into the vertebrate host upon taking a bloodmeal. The sporozoites travel to the liver and multiply asexually within hepatocytes, generating merozoites that are then liberated into the bloodstream, where the intraerythrocytic development cycle (IDC) begins. Repeated rounds of this cycle, which consists of erythrocyte invasion followed by asexual multiplication (including the ring, trophozoite and schizont stages) and release of new merozoites, enables parasites to rapidly increment their biomass and establish an enduring infection in the vertebrate host. However, some parasites convert into sexual forms termed gametocytes and abandon this cycle. These sexual forms sequester in the bone marrow, where they develop for about ten days before returning to the bloodstream as mature stage V gametocytes, which are infective to mosquitoes. Mating and sexual reproduction occur within the mosquito midgut, before replicating asexually again within the mosquito and eventually generating new sporozoites (2).

Life cycle progression in *Plasmodium spp.* is controlled mainly at the transcriptional level, such that each stage is characterized by a specific gene expression programme (3–5). This enables parasites to thrive in the multiple different environments they are exposed to, which entail dramatic differences in conditions such as temperature, pH, nutrient availability or host immune responses. However, in addition to environmental diversity associated with life cycle progression, malaria parasites also have to confront fluctuating conditions within the same niche. These fluctuations, which can occur between individual hosts of the same species or even within the course of a single blood infection, may derive for instance from changes in the host’s physiological or immunological state, or from the effects of antimalarial treatment (6, 7). Adaptation to such diverse conditions occurs through various genetic and non-genetic mechanisms. While in malaria parasites genetic changes play a major role in species evolution and long-term adaptation to new conditions, as in any other organism, rapid adaptation to conditions that fluctuate frequently requires reversible, dynamic mechanisms that provide phenotypic plasticity (i.e., alternative phenotypes from the same genome) (8). One such mechanism is clonally variant gene (CVG) expression, which refers to genes that can be found in a different state (active or silent) in different individual parasites with identical genomes and at the same stage of life cycle progression. While both states are heritable, CVGs undergo low-frequency stochastic switches between the active and silent states, which constantly generates transcriptional heterogeneity within parasite populations (9–11). Changes in the expression of these genes can result in phenotypic variation; therefore, when the conditions of the environment change, natural selection can operate upon this preexisting diversity and eliminate from the population parasites with CVGs expression patterns that do not confer sufficient fitness under the new conditions. This is considered a bet-hedging adaptive strategy (12), a type of adaptive strategy commonly observed in many microbial species (13–15).

All *Plasmodium spp*. appear to have genes under clonally variant expression during asexual blood growth, suggesting that this is a universal feature of malaria parasites. At present, the best characterized CVGs are those from the most virulent human malaria parasite species, *P. falciparum*. These include, among others, large gene families such as *var*, *rif*, *stevor*, *pfmc-2tm*, *hyp*, *phist* and *surfin* linked to pathogenesis, antigenic variation and host cell remodeling; *mspdbl2*, *eba140* and *pfrh4* genes linked to erythrocyte invasion; *clag* genes involved in solute transport; *acs* and *acbp* families linked to acyl-CoA metabolism; and *pfap2-g*, the master regulator of sexual conversion (9–12). Specific adaptive roles have been clearly demonstrated for changes in the expression of *var* (16–18) and *clag3* (19–24) genes, but changes in the expression of other CVGs are likely to also play important roles in adaptation to fluctuating conditions.

The active or silenced state of CVGs is regulated at the epigenetic level (25). These genes are mainly located in subtelomeric bi-stable chromatin domains, in which both the active (euchromatin) and the silenced (facultative heterochromatin) state can be stably transmitted for several generations of asexual growth. Spontaneous transitions between the two states underly the transcriptional switches (8–11). For all CVGs analyzed so far, the silenced state is characterized by the post-translational histone modification histone H3 lysine 9 tri-methylation (H3K9me3) and heterochromatin protein 1 (HP1), whereas the active state is associated with acetylation of H3K9 (H3K9ac). Transmission of these histone modifications through asexual replication constitutes the epigenetic memory for the transcriptional state of CVGs (8-11,26-30). In the *var* and *clag3* families, the only two *P. falciparum* gene families known to show mutually exclusive expression (i.e., only one member of the family is active at a time in individual parasites) (31–35), this epigenetic memory is erased during transmission stages (in the broad sense, including gametocyte, mosquito and liver stages) (23,36–38).

The expression patterns of CVGs under culture conditions have been characterized for several specific gene families and also at a genome-wide scale (12,32,33,39,40). However, understanding how CVGs are expressed under the natural conditions of a human infection is complicated by multiple factors, including common occurrence of polyclonal infections and genetic variation among isolates, which mainly affects CVGs. The expression of only some CVG families such as *var*, *clag3* and genes involved in erythrocyte invasion has been characterized in some detail in natural infections (8,16,23,41–43). This is an important gap of knowledge, because the conditions of the environment affect the expression patterns of CVGs that are selected. Indeed, the limited data available suggest that the transcriptome of malaria parasites grown *in vitro* differs substantially from that observed *in vivo*, including differences in CVGs expression (44, 45).

Parasites obtained from controlled human malaria infection (CHMI) trials provide many of the advantages of both cultured parasites and parasites from natural human infections, because they are exposed to “real” human host conditions but they have a well-defined homogeneous genetic background. Furthermore, the conditions of the host and the time of infection are well-controlled, which reduces the number of variables and facilitates the interpretation of the results. Therefore, CHMI trials are a valuable tool to study malaria parasite biology, especially in relation to their adaptation to different host environments and *in vivo* expression of CVGs. So far, the analysis of gene expression in CHMI samples has focused mainly on the *var* and *clag3* families (23,36–38,46–48). A more recent study analyzed at a genome-wide level the transcriptome of parasites obtained from CHMI volunteers, but only in ring stage parasites, precluding the characterization of expression patterns for genes expressed at other stages of the IDC (49).

To provide a complete view of *P. falciparum* CVGs expression patterns during the initial phase of a blood infection in non-immune human hosts, we performed a genome-wide transcriptomic comparison across the full IDC between parasites obtained from volunteers participating in a CHMI trial and the parental NF54 line. With this controlled approach, we identified transcriptional differences between parasites growing under *in vitro* culture or human circulation conditions, and also tested the hypothesis that CVGs other than the mutually-exclusively expressed *var* and *clag3* genes undergo an epigenetic reset upon passage through transmission stages. To confirm the epigenetic nature of the differences observed, we mapped the genome-wide distribution of heterochromatin under the two different conditions.

## Results

### Passage through transmission stages and exposure to the human host conditions results in changes in CVGs expression

We performed a time-course genome-wide transcriptomic analysis across the full IDC of *P. falciparum* parasites obtained from a CHMI trial in which cryopreserved Sanaria NF54 sporozoites were injected into non-immune (not previously exposed to malaria) human volunteers (50). Parasites were cryopreserved on day 9 after infection and upon microscopy diagnosis on day 11-14. Transcriptomic analysis was performed using samples from four different volunteers (V18, V35, V48 and V63, hereafter collectively termed vNF54) collected on the day of diagnosis. Parasites were thawed and cultured for the minimum number of cycles needed to obtain sufficient material (typically 4 replication cycles) and then tightly synchronized before harvesting RNA at defined time points of the IDC [10-15, 20-25, 30-35 and 40-45 h post-invasion (hpi)]. In parallel, we obtained RNA at the same time points from two independent biological replicates of tightly synchronized cultures of the parental (premosquito) NF54 line (pNF54) (Fig. 1A). Transcript levels were determined using two-channel long oligonucleotide microarrays in which samples where hybridized against a common reference pool to obtain relative expression values (Cy5/Cy3). To quantify transcript level differences between vNF54 and pNF54 lines, we calculated for each gene the maximum average fold change among overlapping time intervals of half the duration of the IDC (mAFC) (12) (see Methods).

**Figure 1.**
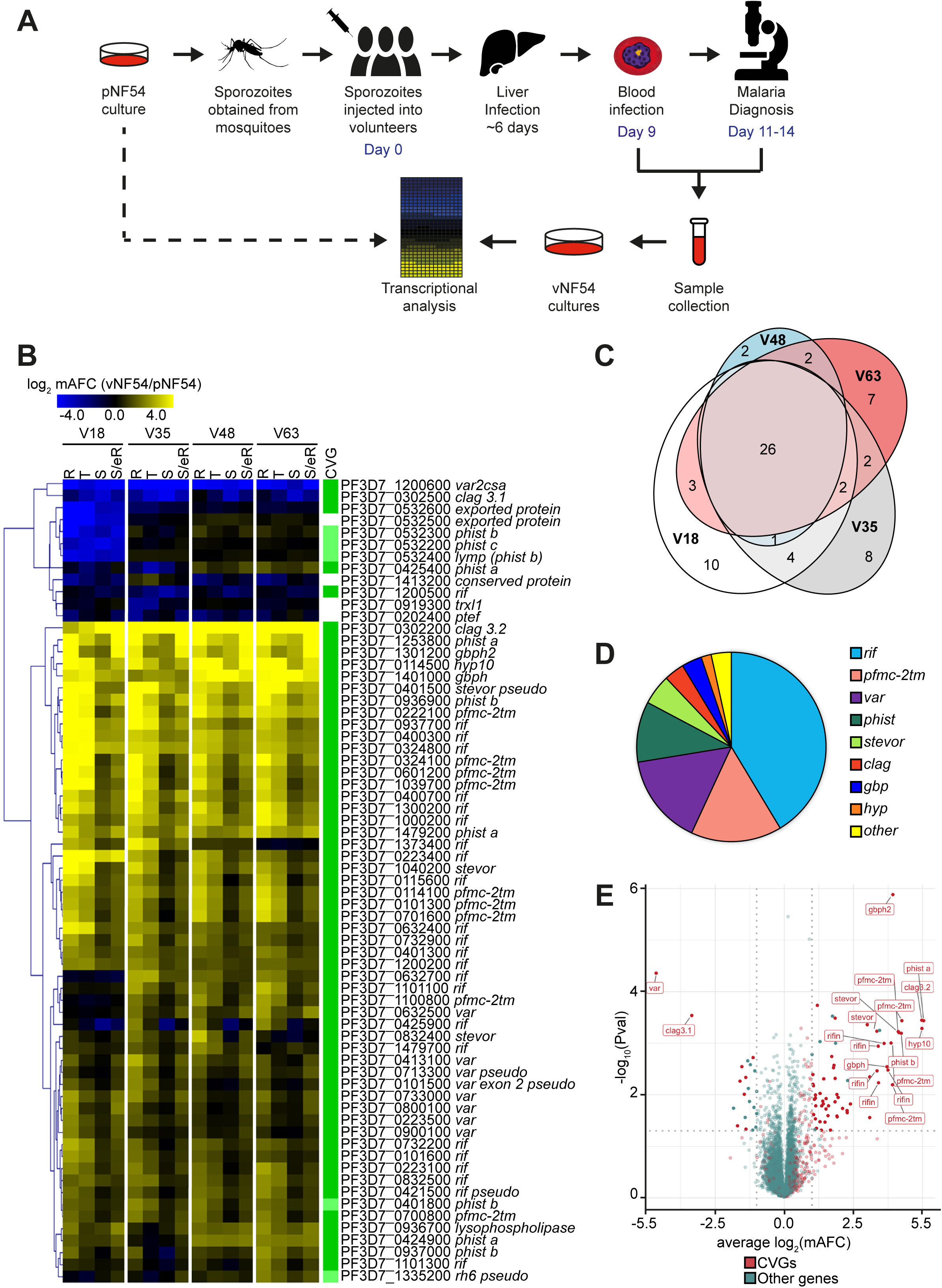
Genes differentially expressed between vNF54 and pNF54 lines. **(A)** Schematic of the study design, comparing transcript levels between vNF54 lines obtained from volunteers participating in a controlled human malaria infection (CHMI) trial and the parental (premosquito) pNF54 line. **(B)** Heatmap of genes differentially expressed between vNF54 and pNF54 lines, ordered by hierarchical clustering. V18-V63 are vNF54 lines obtained from different volunteers. Values are the log_2_ of the average fold-change (AFC) of vNF54 vs pNF54 lines over four overlapping time intervals corresponding to the stages indicated (R: rings, T: trophozoites, S: schizonts, S/eR: schizonts and early rings). Only genes with an absolute value of the log_2_ of the maximum AFC (mAFC) >2 (relative to pNF54) in at least one of the vNF54 lines are shown. The column at the right indicates whether a gene was previously classified as a CVG (dark green, see Supplementary Table 2) or belongs to a gene family in which other genes are CVGs (light green). **(C)** Euler diagram showing the overlap between genes with an absolute value of the log_2_(mAFC) >2 between the different volunteers. **(D)** Pie chart showing the distribution of genes with an absolute value of the log_2_(mAFC) >2 between different gene families. **(E)** Volcano plot representing expression differences between vNF54 and pNF54 lines. Expression fold-change values for each gene are the average of the log_2_(mAFC) in the four vNF54 lines. *P*-values were calculated using an unpaired two-sided *t*-test. CVGs are shown in red and other genes in green. Name labels are provided for genes with a mAFC >10 and a *P*-value <0.01. Vertical dotted lines mark a log_2_(mAFC)=1 or −1, whereas the horizontal dotted line marks a *P* value=0.05.

There was a total of 67 genes with a mAFC >4 in either direction [absolute value of the log_2_(mAFC) >2] between at least one of the vNF54 lines and the pNF54 line, 21 of which had a mAFC >16 (Fig. 1B and Supplementary Table 1). The vast majority of differentially expressed genes had been previously identified as CVGs (Fig. 1B), either based on differential expression between isogenic lines (12) or because they carried the H3K9me3 or HP1 heterochromatin marks (62–65) (Supplementary Table 2). Overall, the transcriptional changes (relative to pNF54) observed in the four vNF54 lines were almost identical (Fig. 1B), with the exception of a few genes showing a different pattern only in the V18 line. As a consequence, there was a large overlap in the genes differentially expressed (relative to pNF54) between each of the four vNF54 lines (Fig.1C). Closer analysis revealed only a small number of genes with large transcript level differences among the four vNF54 lines, and many of them showed a different pattern only in the V18 line, including a cluster of five downregulated neighbor genes in chromosome 5 (Supplementary Fig. 1A).

The majority of genes differentially expressed in all vNF54 lines compared to pNF54 were expressed at higher levels in vNF54 lines, but a small number of genes were expressed at lower levels. The two most downregulated genes (mAFC >10) were *var2csa* and *clag3.1*. The most upregulated genes in vNF54 lines (mAFC >32) were a *hyp10* gene encoding an exported protein (PF3D7_0114500) (66), a *phist-a* gene (PF3D7_1253800), *clag3.2* (PF3D7_0302200) and a *pfmc-2tm* gene (PF3D7_0324100). Overall, the majority of genes that showed changes in expression between pNF54 and vNF54 lines belong to the large *pfmc-2tm*, *rif*, *var*, *stevor* and *phist* CVG families encoding exported proteins (16-18,66,67), or to the smaller families *clag*, involved in solute transport (20), and *gbp*, of unknown function (68) (Fig. 1D). Given the overall similarity between the different vNF54 lines, we performed an additional analysis in which the different vNF54 lines were treated as replicates, which enables statistical analysis of the expression differences observed. The genes identified as most differentially expressed in this analysis were roughly the same as in the analysis based on each vNF54 line treated individually, and *gpbh2* (PF3D7_1301200) was the most significantly upregulated gene in vNF54 lines (Fig.1E and Supplementary Table 1).

### Transcriptional changes in specific CVG families

To analyze the expression patterns of CVGs within specific gene families, in addition to the transcript levels relative to a common reference pool (Cy5/Cy3 values) we also used the sample signal (Cy5 channel). The two-channel microarray approach used here is designed for hybridization against a common reference pool, which enables robust comparison of relative transcript levels between samples (3,12,52). However, the sample signal alone is informative as a semi-quantitative estimate of the expression intensity of the genes, because it enables the identification of the predominantly expressed members of specific families (12), rather than informing only on the relative levels between samples.

Analysis of the transcriptional differences between pNF54 and vNF54 lines for the specific CVG families to which the majority of differentially expressed genes belong revealed different scenarios for different families (Fig. 2A-B). Essentially all members of the *pfmc-2tm* family (39,67,69) were strongly upregulated in vNF54 lines (mAFC > 8 in all but three genes) (Fig. 2A,C), and this was confirmed analyzing the Cy5 signal only, which revealed a large increase in the expression of the majority of *pfmc-2tm* genes (Supplementary Fig. 2A). Protein levels for a specific PfMC-2TM protein for which antibodies were available also revealed clearly increased expression in a vNF54 line compared to pNF54 (Supplementary Fig. 2B). In contrast, in the *var*, *rif*, *phist* and *stevor* families several specific genes were upregulated in vNF54 lines, but many other genes did not change and, in some families, a few were downregulated. In the case of the mutually exclusively expressed *var* genes, vNF54 lines showed upregulation of many *var* genes, especially of type B, whereas the type E *var2csa* gene was strongly downregulated (Fig. 2B-C). Analysis of the Cy5 values revealed that *var2csa* was the predominantly expressed *var* gene in pNF54, whereas in vNF54 lines multiple *var* genes were expressed at intermediate levels, similar to findings from previous CHMI studies (36,38,46) (Supplementary Fig. 2C). Considering that individual parasites express a single *var* gene, this result indicates that the parental population was relatively homogeneous, such that the vast majority of individual parasites expressed the same *var* gene (*var2csa*). In contrast, after passage through transmission stages the population became heterogeneous, with different individual parasites expressing different *var* genes.

**Figure 2.**
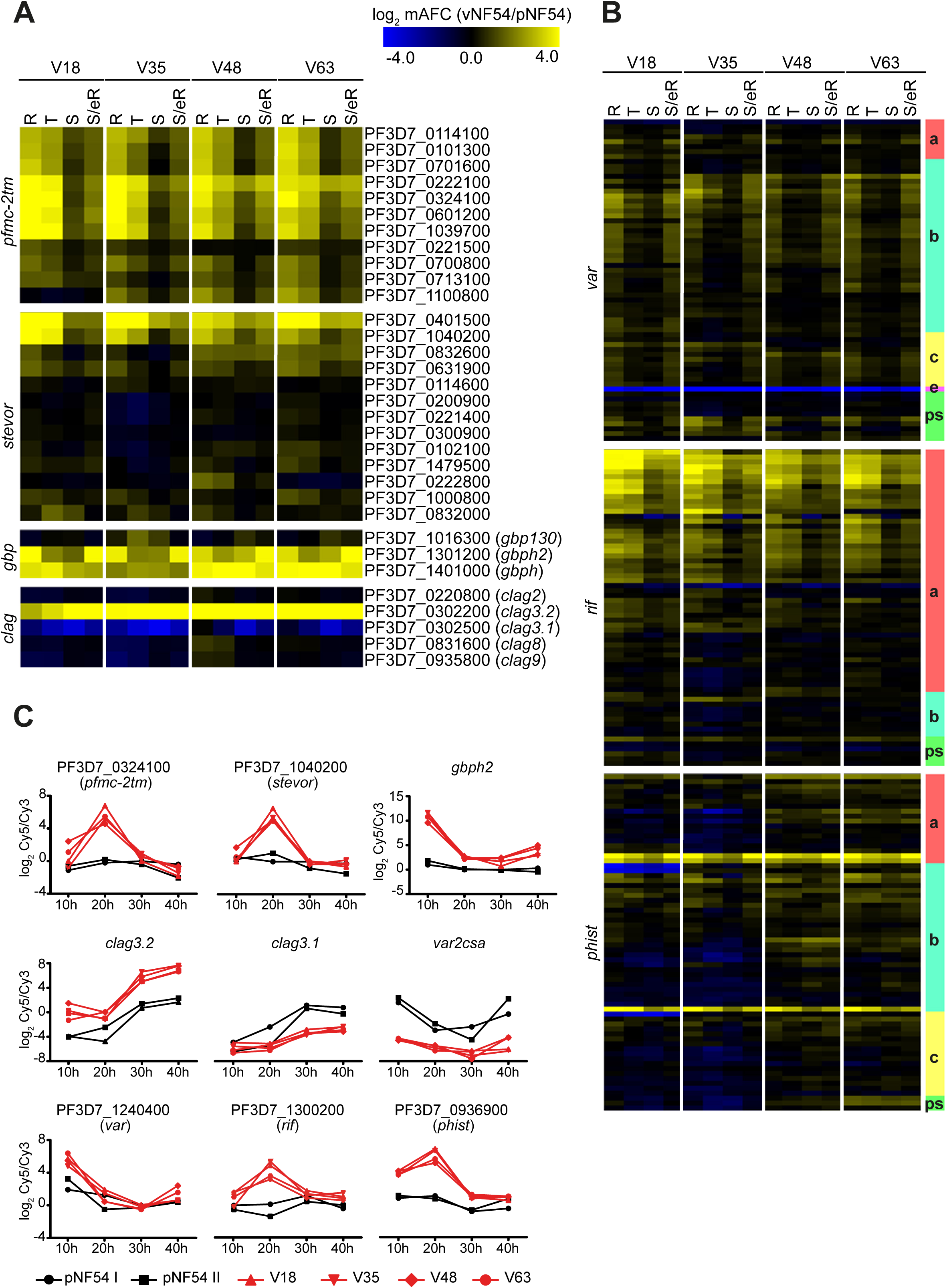
Transcriptional differences in specific CVG families. **(A-B)** Transcript level changes in vNF54 lines relative to pNF54 lines, as in Fig. 1B, for all genes analyzed of the CVG families indicated. In panel **B**, the subfamily of each gene (as annotated in PlasmoDB or in a previous study (12)) and pseudogenes (ps) are indicated in the column at the right. **(C)** Time course expression plots of selected genes of different CVG families. Values are the log_2_ of the sample vs reference pool ratio (Cy5/Cy3) in the vNF54 and pNF54 lines.

In the *rif* family (18,70–73), a large subset of genes encoding type A RIFINS were upregulated in all vNF54 lines, whereas no general differences were observed in genes encoding type B RIFINS. In the *stevor* and *phist* families (18,66,74) only a small subset of specific genes showed strong upregulation in vNF54 lines, essentially the same genes in all the four lines (Fig. 2A-C). In the *rif*, *stevor* and *phist* families, which participate in antigenic variation and show expression switches but do not show mutually exclusive expression (39,73,75), the analysis of the Cy5 signal revealed that in spite of changes in the expression of some specific genes, the predominantly expressed genes are similar between pNF54 and vNF54 lines (Supplementary Fig. 2D-F). None of the genes that showed increased transcript levels in vNF54 lines became the dominantly expressed gene in any of these families.

Two small CVG families, *gbp* (68) and *clag* (20), showed major changes in the expression of a large proportion of their genes. Of note, two of the three members of the *gbp* family, *gbph* and *gbph2*, were among the most strongly upregulated genes (mAFC >16) in vNF54 lines (Fig. 2A,C and Supplementary Fig. 2G). In all vNF54 lines, there was increased expression of *clag3.2* and reduced expression of *clag3.1*, consistent with our previous RT-qPCR results showing that in the pNF54 population essentially all parasites express *clag3.1*, whereas in vNF54 lines there is a mixture of parasites expressing *clag3.1* and parasites expressing *clag3.2* (23). There was no major change in the expression of the other CVG of the *clag* family, *clag2* (32) (Fig. 2A,C and Supplementary Fig. 2H).

In contrast to the major expression changes observed in the gene families described above, the expression of other CVG families such as acyl-CoA synthetases (*acs*), acyl-CoA binding proteins (*acbp*), *lysophospholipases*, exported protein kinases (*fikk*), *hyp*, *surfin* and families linked to erythrocyte invasion (*eba*, *pfrh*) was almost identical between pNF54 and vNF54 lines, with the exception of specific genes such as *hyp10* (PF3D7_0114500) (Supplementary Fig. 3).

### Large changes in CVGs expression between vNF54 and pNF54 are determined at the epigenetic level

The observation that the majority of genes differentially expressed between pNF54 and vNF54 lines are CVGs, which have been previously shown to be regulated by truly epigenetic mechanisms (25), strongly suggests that parasites recovered from the volunteers mainly differ from the parental line in their epigenetic make up, rather than at the genetic level. To confirm this view, we sequenced the whole genome of the pNF54 and two vNF54 lines (V18 and V63) and found no genetic differences in either coding or non-coding regions that are likely to explain the transcriptional changes common to all vNF54 lines (Supplementary Table 3). However, we observed several large subtelomeric deletions in V18 that explain the reduced expression of several genes specifically in this parasite line, including the cluster of neighbor genes in the subtelomeric regions of chromosome 5 (Supplementary Table 3 and Supplementary Fig. 4).

Next, we performed comparative H3K9me3 chromatin immunoprecipitation followed by sequencing (ChIP-seq) analysis to determine the distribution of heterochromatin in pNF54 and one of the vNF54 lines (V63). In the genes showing the largest transcript level differences between the two lines, higher expression was associated with absence of heterochromatin in the upstream regulatory regions, whereas lower expression was associated with presence of heterochromatin (Fig. 3A,C and Supplementary Table 4). However, there were no apparent differences in heterochromatin prevalence between V63 and pNF54 in many genes from large CVG families that were expressed at higher levels in V63, such as *pfmc-2tm*, *rif* and *var* genes (Fig. 3B,C). The most plausible interpretation for this result is that these genes were activated in only a small subset of the parasites in vNF54, accounting for the transcript levels increase observed, but they remained silenced and in a heterochromatic state in the majority of parasites. In the case of *var* genes, this interpretation fits with the know mutually-exclusively expression of this family.

**Figure 3.**
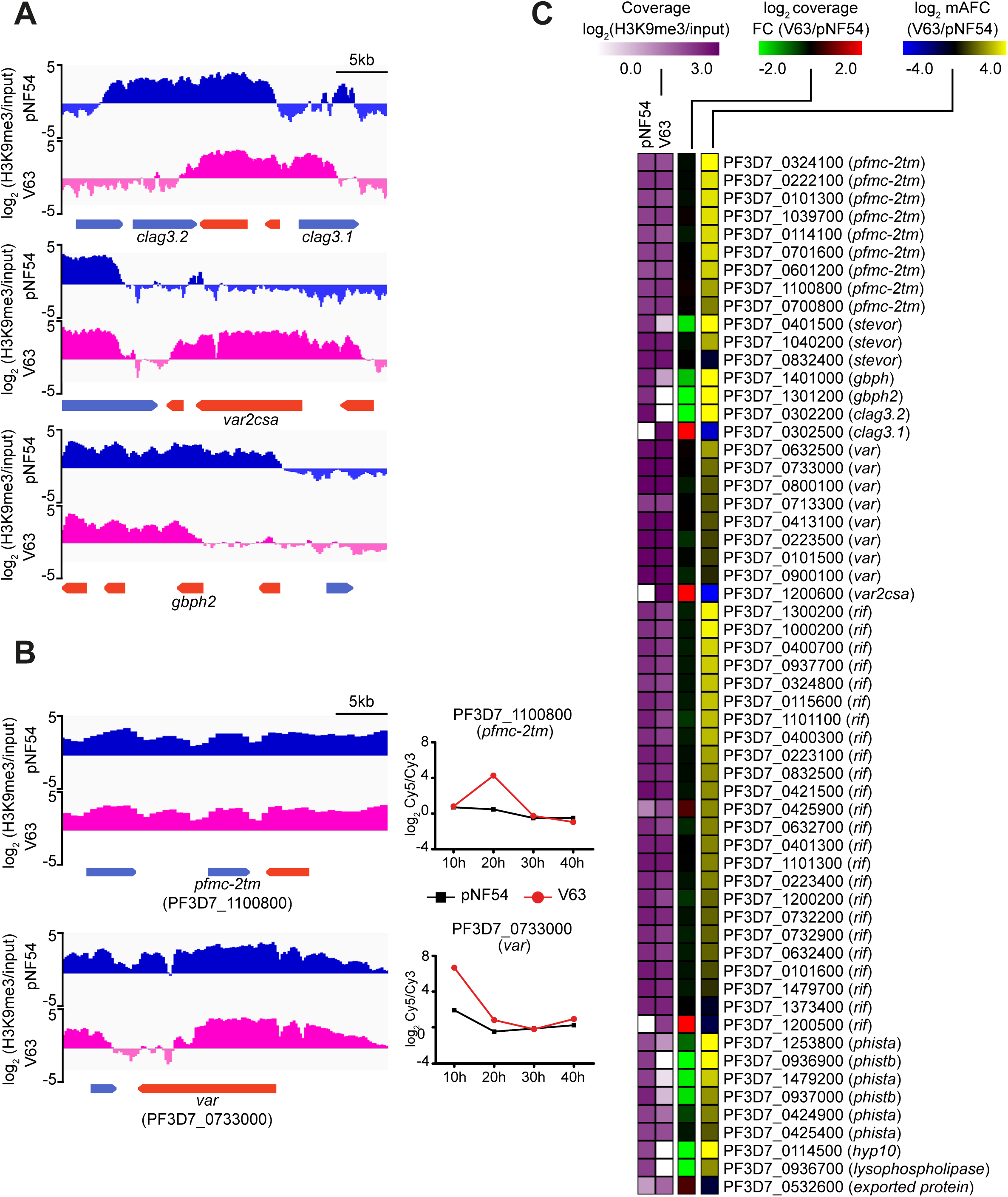
ChIP-seq analysis of vNF54 and pNF54 lines. **(A-B)** Distribution of normalized H3K9me3 signal relative to input at selected loci in the pNF54 line and the vNF54 line V63. The time-course expression plot for these two lines is shown for genes not included in Fig. 2C. **(C)** H3K9me3 ChIP-seq coverage (from −1,000 bp or closest upstream gene to +500 bp, from the ATG), coverage fold-change (FC) between V63 and pNF54, and mAFC between V63 and pNF54 for the same genes displayed in Fig. 1B, except for non-CVGs (which do not have heterochromatin).

### Changes in the expression of CVGs are generally determined by a reset of epigenetic patterns during transmission stages

For *var* and *clag3* genes, an “epigenetic reset” during transmission stages results in a broad pattern of expression in the parasites population at the onset of a blood infection, followed by progressive selection of parasites expressing variants that confer higher fitness under the conditions of the human host circulation (23,36–38,46,48). To determine if the epigenetic memory for the transcriptional state of other CVGs is also lost during transmission stages, we compared the expression of *gbph2*, *pfmc-2tm* and *stevor* genes between samples collected from the volunteers on day 9 or on the day of microscopy diagnosis (for the samples used, day 11 or 14). Since liver stage development lasts ∼6 days (36,46,47,76), day 9 samples multiplied in the human circulation for only one round of the IDC after egress from the liver (second generation blood stages), whereas day 11 / 14 samples replicated for 1 or 3 additional cycles, respectively (third or fifth generation blood stages), according to previous estimations (47). We reasoned that if within-host selection alone was responsible for the differences between pNF54 and vNF54 lines, day 9 samples would show expression levels intermediate between pNF54 and day 11 / 14 samples, as parasites with unfavorable expression patterns would be eliminated progressively. In contrast, if an epigenetic reset is the main determinant of the expression patterns observed, similar transcript levels would be expected between day 9 and 11 / 14 samples. This same rationale was previously used to conclude that the expression patterns of *clag3* genes are reset during transmission stages (23).

Because the parasitaemia of day 9 samples was very low, obtaining sufficient material for RT-qPCR analysis required ∼7 cycles of *in vitro* growth, whereas only ∼4 cycles were needed for day 11 / 14 samples. Therefore, we first determined whether extended culture affected the expression levels of these genes. After five weeks in culture, the expression levels of *gbph2*, one *pfmc-2tm* gene, one *stevor* gene, and total *pfmc-2tm* family transcripts remained stable in both pNF54 and a vNF54 line (V63). This result indicates that growth under *in vitro* conditions does not rapidly alter the expression of these genes and does not represent a confounding factor for the comparison of day 9 with day 11 / 14 samples (Fig. 4A). These experiments also confirmed the much higher transcript levels for these genes in vNF54 than in pNF54, and revealed that they are virtually silenced in the latter.

**Figure 4.**
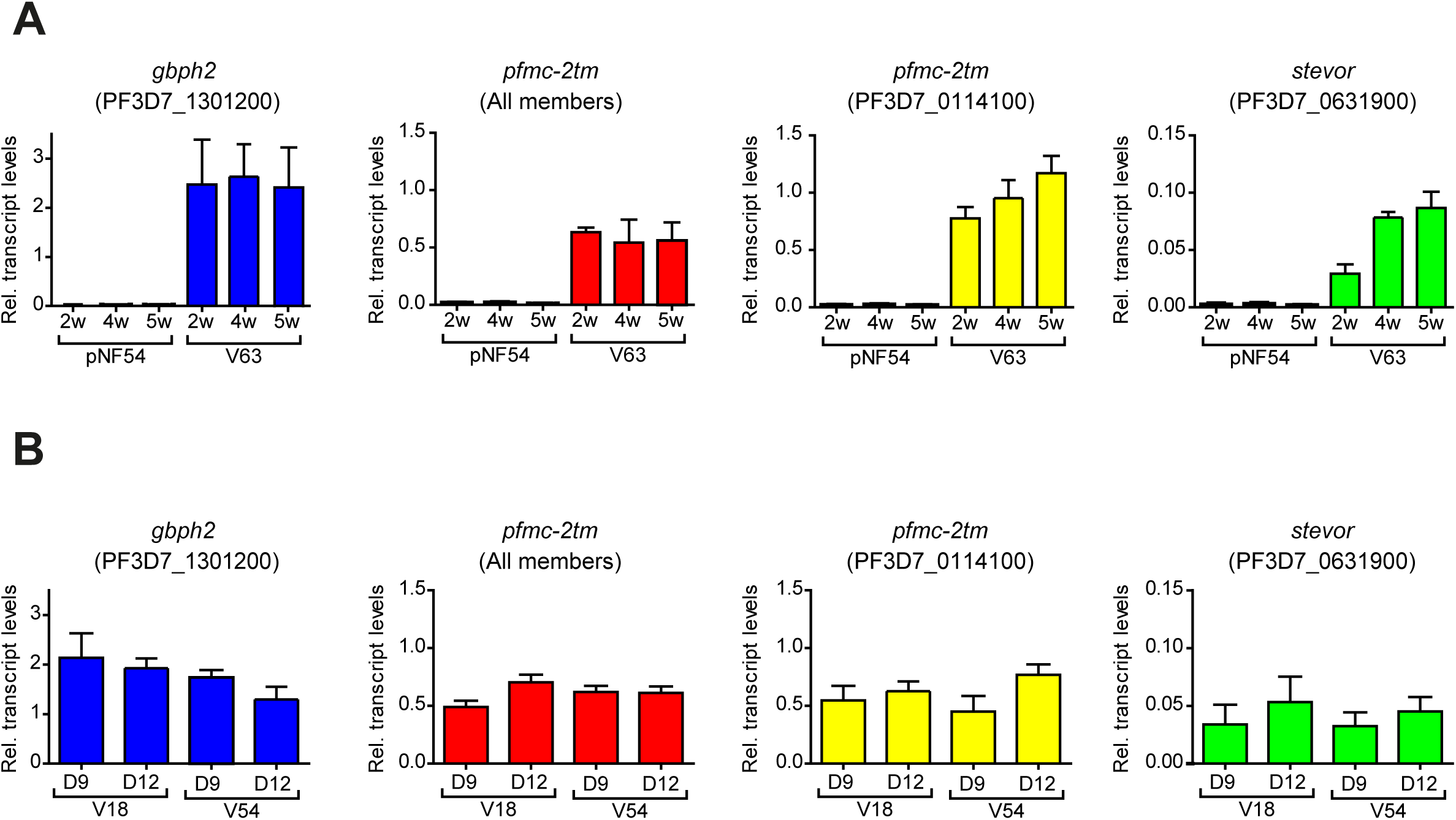
Changes in transcript levels associated with time in culture or time in the human circulation. **(A)** Effect of duration of growth under culture conditions on the expression of selected CVGs with different transcript levels between V63 (a vNF54 line) and pNF54. Relative transcript levels were determined by RT-qPCR after different times in culture, up to 5 weeks (w), in the pNF54 and V63 lines. RNA for transcriptional analysis was collected at the ring stage (10-15 h post-invasion, hpi) for the *gbph2* gene, and at the early trophozoite stage (20-25 hpi) for *pfmc-2tm* and *stevor* genes. For “*pfmc-2tm* (All members)”, primers that amplify all genes of the family were used. Transcript levels are normalized against *serine-tRNA ligase (serrs)*. Data are presented as the average and s.e.m. of three independent biological replicates. **(B)** Comparison of relative transcript levels between parasites collected at day 9 post-infection or at days 14 (V18) or 11 (V54) (day of microscopy diagnosis), determined as in the previous panel. No significant differences (*P*<0.05) were observed between different times in culture (panel A) or between day 9 and day 11 / 14 samples (panel B) using two-sided *t*-tests.

Next, we compared the expression levels of these genes between parasites collected on day 9 and on the day of microscopy diagnosis in two vNF54 lines (V18 and V54, with microscopy diagnosis on day 14 and 11, respectively), which revealed no major differences (Fig. 4B). Therefore, these results are consistent with the idea that there is a general epigenetic reset of CVG expression patterns during passage through transmission stages, beyond the mutually exclusively expressed *var* and *clag3* families.

### Phenotypic comparison of pNF54 and vNF54 lines

We performed exploratory experiments to determine if the transcriptional differences observed between pNF54 and vNF54 lines result in measurable functional differences. Since many of the genes showing differential expression are exported to the erythrocyte cytoplasm or membrane (18,66,67) and some directly impact the mechanical properties of infected erythrocytes (i.e., *stevor*) (77), we compared the membrane deformability at the trophozoite stage of pNF54- and vNF54 (V63)-infected erythrocytes using a microsphiltration assay. While V63 showed a higher retention rate, indicative of lower deformability, the difference was not significant (Fig. 5A).

**Figure 5.**
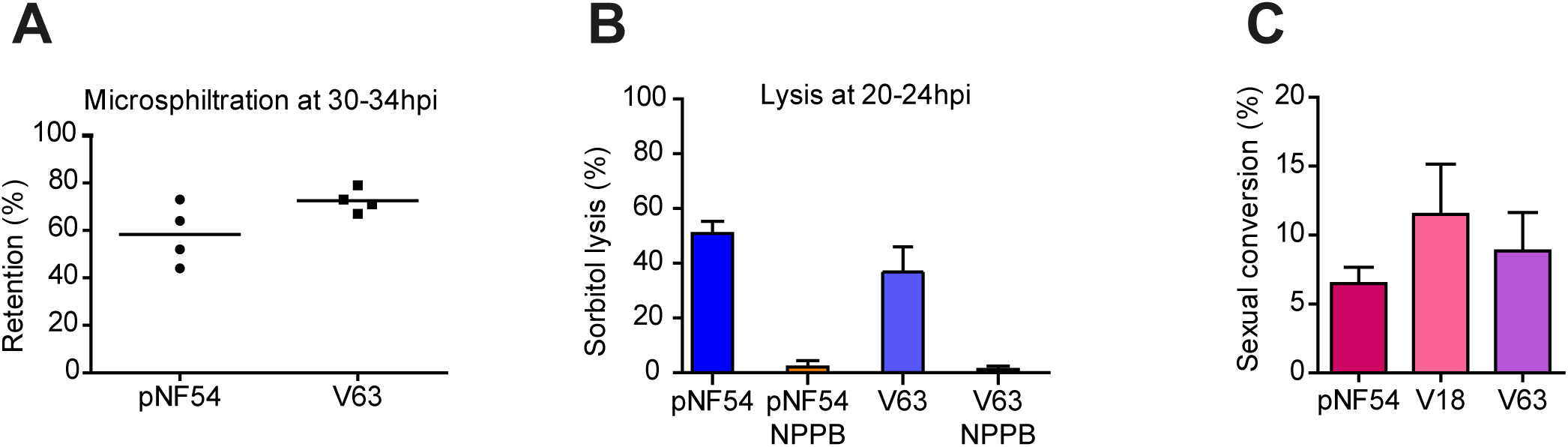
Phenotypic comparison of vNF54 and pNF54 lines. **(A)** Retention in microsphiltration assays of pNF54 and a vNF54 line (V63). Data are presented as the average and s.e.m. of four independent biological replicates. **(B)** Sorbitol lysis assays for pNF54 and V63 in the absence or presence of NPPB (an inhibitor of new permeation pathways). Data is presented as the average and s.e.m of three independent biological replicates. **(C)** Sexual conversion rate (proportion of parasites that convert into sexual forms) of pNF54 and two vNF54 lines (V18 and V63). Data are presented as the average and s.e.m. of three independent biological replicates. No significant difference (*P*<0.05) was observed between pNF54 and vNF54 lines in any of the panels, using two-sided *t*-tests.

Since we observed changes in the expression of *clag3* genes, which determine solute uptake in infected erythrocytes (20), we compared sorbitol permeability between pNF54 and vNF54 lines. Sorbitol uptake resulting in red cell lysis was observed at similar levels in both lines (Fig. 5B), as expected given that only the simultaneous silencing of the two *clag3* genes, which did not occur in either pNF54 or vNF54, is known to prevent sorbitol uptake (22). Lastly, we compared the sexual conversion rates of pNF54 and vNF54 lines (V18 and V63), which revealed no significant differences (Fig 5C). This is consistent with the similar expression levels of *pfap2-g*, the master regulator of sexual conversion (78), and early gametocyte markers between pNF54 and vNF54 lines.

## Discussion

Here we provide the first genome-wide transcriptional characterization across the full IDC of *P. falciparum* in the initial days of a blood infection in non-immune humans, and compare it with the transcriptome of parasites with the same genome but maintained under *in vitro* culture conditions. We also compared the genome-wide distribution of heterochromatin between parasites obtained from the infected humans or in culture. We found that the largest expression differences occur in CVGs as a consequence of epigenetic changes in the distribution of heterochromatin, and provide an accurate view of how parasites use their CVGs when they establish a new blood infection after egress from the liver. We also show that passage through transmission stages results in a general reset of the epigenetic memory involving many CVG families, but the ensuing alterations in the patterns of expressed and silenced genes differ between gene families. Thus, the expression of CVGs *in vivo* is governed by low frequency switches and selective pressures during asexual growth (9–11), and also by a reset during transmission stages.

Several studies comparing gene expression patterns between parasites obtained from naturally infected patients and non-matched culture adapted lines reported higher expression *in vivo* for many CVGs, especially those encoding erythrocyte surface antigens (8,44,45,72,79–84). This is concordant with our observation that there were many more genes showing higher expression in parasites recovered from the volunteers than vice versa, suggesting that long-term growth in culture results in progressive silencing of many CVGs. However, in some gene families this pattern can also be explained by increased transcriptional heterogeneity in parasite populations obtained from the volunteers, as shown to be the case for *var* or *clag3* genes (23, 37). The expression pattern of some specific genes that in our dataset were expressed at higher levels in parasites from the volunteers, such as *gbph2* or *clag3.2*, is also consistent with previous reports of higher expression of these genes in natural infections compared to cultured parasites (23, 83).

The majority of changes between premosquito parasites and parasites obtained from the infected volunteers occurred in gene families involved in processes such as antigenic variation, erythrocyte remodeling or solute transport, whereas other CVG families such as *acs*, *acbp*, *fikk*, *surfin*, *eba* or *pfrh* showed few alterations. Even within the same family, subgroups of genes with different predicted functions showed different expression patterns: type A RIFINS, predicted to encode proteins localized in the infected erythrocyte surface and involved in rosetting (18, 73), were generally upregulated in parasites collected from the volunteers, but not type B RIFINS with different predicted function and localization. The most dramatic changes were observed in the *pfmc-2tm*, *gbp*, *var* and *clag3* families, whereas other families such as *rif*, *stevor* or *phist* showed important transcript level differences for several specific genes but the predominantly expressed genes and global expression levels of the family were not altered. Together, these observations suggest that some CVG families and subfamilies have high plasticity in their expression patterns, whereas others have clearly “preferred” patterns even under different environment conditions.

While the epigenetic state of CVGs is transmitted from one generation of asexual parasites to the next (11,17,25), it has been proposed that an “epigenetic reset” may occur during transmission stages, such that the epigenetic memory for the transcriptional state of all CVGs is erased and new patterns of expressed and silenced CVGs are reestablished stochastically (85). In support of this idea, in patients infected with *Plasmodium* spp. to treat neurosyphilis, malaria severity was higher when infection was transmitted by serial blood passage rather than by mosquito bite. This suggests that repeated cycles of the IDC progressively select for parasites with CVGs expression patterns associated with increased fitness (which may result in higher virulence when changing host by blood passage, but in long-term, chronic infection in the same host is counteracted by acquired immune responses), whereas passage through transmission stages results in a reset of the epigenetic patterns and lower virulence (85). Experimental support for this hypothesis was obtained in the murine malaria parasite *P. chabaudi*, as mosquito-transmitted parasites were less virulent than those transmitted by blood passage, and expressed a wider repertoire of CVGs (85, 86). Evidence for an epigenetic reset during transmission stages in *P. falciparum* was previously obtained for *var* and *clag3* genes, as parasite populations recovered from volunteers participating in CHMI trials showed expression patterns for these genes clearly distinct from the parental population, with a broad expression pattern that was indicative of heterogeneity between individual parasites (23,36–38). Here we observed clearly distinct expression patterns between premosquito parasites and parasites obtained from the volunteers for multiple gene families, and similar expression patterns between day 9 and day 11-14 samples in all the CVG families analyzed (*gbp*, *pfmc-2tm* and *stevor*). Therefore, our results are consistent with a general epigenetic reset of CVGs during transmission stages, beyond genes showing mutually exclusive expression. The postulated general reset results in high population heterogeneity for the expression of many CVGs and consequently a phenotypically and antigenically diverse population when parasites exit the liver, which favors survival of the population in the unpredictable conditions of the blood of a new human host (a bet-hedging survival strategy). Later on, the conditions of each particular human host exert selective pressure and natural selection shapes the epigenetic patterns that will prevail during continued blood growth, as previously shown specifically for *var* genes (36,38,48). However, the long-term progressive selection of subsets of parasites with specific patterns of expressed and silenced CVGs could not be monitored in our CHMI trial, because the infection was terminated as soon as parasites were detected by light microscopy or symptoms appeared.

We postulate that during transmission stages the epigenetic memory is erased and new epigenetic patterns are established stochastically, such that they can differ among individual parasites with the same genome. However, the probability of each gene being in an active or silenced state is hardwired and dictated by the underlying DNA sequence. This probabilistic scenario can explain why in some gene families the expression state of the majority of genes was identical between pNF54 and vNF54 lines, as in these families this particular state would be greatly favored during the *de novo* establishment of epigenetic patterns. Of note, in some genes from specific families, we observed large transcript level fold-differences that were not accompanied by changes in the levels of H3K9me3 occupancy. One such family is *var*, which shows mutually exclusive expression. We observed increased transcript levels of many *var* genes and reduced expression of a single *var* gene in the parasite population obtained from the volunteers, similar to previous CHMI studies (36,37,46). This pattern is explained by a shift from a homogeneous parental population in which the majority of parasites express the same *var* gene (*var2csa*) to a heterogeneous population in which different individual parasites express a different *var* gene. Therefore, in the parasite populations obtained from the volunteers each individual gene is active in only a small subset of the parasites (e.g. 1%), which still can result in a large fold-increase in transcript levels if fewer parasites expressed it in the premosquito population (e.g., ten-fold increase if it was active in only 0.1% of the premosquito parasites). However, this would result in a neglectable difference in the levels of the heterochromatin mark (e.g., 99.9% vs. 99% of the parasites carry heterochromatin at the locus from the example above), which is not detectable in a comparative ChIP-seq analysis. We propose that a similar situation occurs for *pfmc-2tm*, *rif* and *stevor* genes that show increased transcript levels but no difference in H3K9me3 levels in parasites from the volunteers. This would imply that these genes were activated in only small subsets of parasites. In support of this view, the majority of these genes were not among the predominantly expressed genes within their family. Of note, these gene families do not show strict mutually exclusive expression, but do show variant expression and a limited number of genes of each family are expressed at a time in individual parasites (39,73,75). Future research using single cell approaches (5, 87) will be needed to gain definitive evidence for the model proposed here.

Except for several deletions occurring in parasites recovered from one particular volunteer (V18), parasites recovered from different volunteers were almost identical at both the transcriptomic and genetic levels. If selection of epigenetic variants plays a role in shaping the patterns of CVGs expression in the few cycles of blood growth before sample collection, the selective pressures encountered by the parasites are expected to be the same between different volunteers, because all of them were healthy adults not previously exposed to malaria. Furthermore, if *de novo* establishment of heterochromatin patterns at CVGs after an epigenetic reset is a probabilistic event dictated by the underlying DNA sequence, as we hypothesize, it is also expected to result in similar patterns between parasites in different human individuals. Therefore, it is not surprising that the transcriptome of the parasites from the four different volunteers was almost identical.

Altogether, a model emerges in which a blood infection in a new host starts with a transcriptionally heterogeneous parasite population, consisting of individual parasites with different combinations of active and silenced CVGs from several gene families. Our data provides an accurate picture of how parasites use their CVGs in this initial phase of a blood infection, revealing that they generally express at higher levels than cultured parasites the majority of genes of the *pfmc-2tm* and *gbp* families, as well as specific genes of the *rif* (type A), *phist*, *stevor* and other CVG families. Besides, different individual parasites express different members of the mutually exclusively expressed *var* and *clag3* families, showing a broad expression pattern at the population level (23, 37). During continuous growth in the human blood, subpopulations of parasites with CVGs expression patterns that confer high fitness under the metabolic, drug treatment and immune conditions of a specific human host are progressively selected, whereas others are eliminated. Low frequency switches ensure that new epigenetic variants are constantly generated to ensure population survival when the conditions of the host fluctuate, e.g. by developing acquired immunity. Further studies will be needed to fully characterize the precise phenotypic and antigenic differences that result from specific changes in CVGs expression.

## Materials and Methods

### Human samples and parasite culture

The *P. falciparum* NF54 line at Sanaria (pNF54) and lines derived from human volunteers participating in a CHMI study with Sanaria NF54 sporozoites (vNF54 lines) (50) were cultured in B+ erythrocytes at 3% hematocrit in RPMI-1640-based culture medium supplemented with 10% human serum, under continuous shaking at 100 rpm and in a 5% CO_2_, 3% O_2_ and 92% N_2_ atmosphere.

### Transcriptomic analysis using microarrays

Time-course transcriptomic analysis was performed using cultures tightly synchronized to a 5 h age window, which was achieved by percoll purification of schizonts followed by sorbitol lysis 5 h later (51). Cultures were split into four flasks that were cultured undisturbed for different periods of time (10, 20, 30 or 40 h) before collection in TRIzol and freezing at - 80°C.

RNA was purified using the TRIzol method and cDNA was synthesized by reverse transcription, purified and labeled as previously described (52). Samples were analyzed using two-color long oligonucleotide-based custom Agilent microarrays. The microarray design was based on Agilent design AMADID #037237 (52), modified by adding new probes for genes lacking unique probes and for some ncRNAs and reporter genes (new designs: AMADID-084561 and AMADID-085763) (53). All samples, labeled with Cy5, were hybridized against a common reference pool labeled with Cy3, consisting of a mixture of equal amounts of cDNA from rings, trophozoites and schizonts of pNF54. Microarray hybridization was performed as previously described (53). Images were acquired using a Microarray Scanner (no. G2505C, Agilent Technologies) located in a low ozone hood.

### Microarray data analysis

Initial processing of microarray data, including linear and LOWESS normalization, was performed using Feature Extraction software (Agilent), with default options. The next steps of the analysis were performed using Bioconductor in an R environment (R version 3.5.3). For each individual sample and channel (Cy3 and Cy5), background signal was calculated as the median of the 100 lowest signal probes. Probes with both Cy3 and Cy5 signals below three times the array background were excluded from further analysis. Gene level log_2_(Cy5/Cy3) values and statistical estimation of parasite age (54) were computed as previously described (12). For each gene, the log_2_(Cy5/Cy3) values were plotted against the statistically-estimated culture age (in hpi) and the plots were divided in four overlapping time intervals of identical length that roughly corresponded to the rings, trophozoite, schizont and late schizont/early ring stages (12). For each gene and time interval, the average expression fold-change (AFC) between each vNF54 and its control pNF54 line (the replicate of pNF54 analyzed in parallel) was calculated from the difference in the area under the curve in the log_2_(Cy5/Cy3) vs age plots. The maximum AFC (mAFC) was the value of the AFC in the time interval at which it had the highest absolute value. Visual inspection was used to exclude genes with apparent artefacts from further analysis (e.g. genes with large expression differences observed only at time intervals that don’t correspond to their peak expression). Genes with expression intensity (Cy5 channel) values within the lowest 20^th^ percentile were excluded from the analysis of expression fold-changes between vNF54 and pNF54 lines because differences in genes expressed at near background levels are of low confidence. Heatmaps and hierarchical clustering based on Euclidean distance were generated using TMEV 4.9 (55). For the generation of volcano plots, samples from the different volunteers were treated as replicates and an unpaired two-sided t-test was performed between volunteer and parental NF54 replicates.

### RT-qPCR transcriptional analysis

RNA was purified from parasite samples collected in TRIzol (Invitrogen) using the RNeasy Mini Kit (Qiagen) as previously described (23). Next, purified RNA was reverse transcribed using the Reverse Transcription System (Promega) alongside parallel reactions without reverse transcriptase to exclude gDNA contamination. Quantitative PCR to analyze cDNAs was performed in triplicate wells using the Power SYBR Green Master Mix (Applied Biosystems) in a StepOnePlus Real Time PCR System, essentially as previously described (26, 51). Relative transcript levels were calculated using the standard curve method (51) and the normalizing gene *serine-tRNA ligase* (*serrs*), which shows stable transcript levels across the IDC. The primers used for qPCR are described in Supplementary Table 5.

### Whole genome sequencing and data analysis

PCR-free whole genome Illumina sequencing was used to sequence the whole genome of the parental NF54 line alongside two vNF54 lines (V18 and V63). Genomic DNA was sheared to ∼150-400 bp using a Covaris S220 ultrasonicator and the NEBNext® Ultra™ DNA Library Prep Kit for Illumina was used for library preparation with specific paired-end TruSeq Illumina adaptors for each sample. Due to the high AT content of the *P. falciparum* genome, the End Repair incubation step at 65°C was omitted. After quality control using the Bioanalyzer DNA High Sensitivity kit (Agilent) and quantification using the KAPA Library Quantification Kit (Roche), libraries were sequenced using the Illumina HiSeq2500 System. Over 6 million 125 bp paired-end reads were obtained for each sample.

For the data analysis, read quality was checked (FastQC program) and adaptors were trimmed (Cutadapt program) before mapping sequenced reads to the PlasmodDB *P. falciparum* 3D7 reference genome version 46 (https://plasmodb.org/plasmo/) using the Bowtie2 local alignment algorithm. Next, GATK-UnifiedGenotyper was used to perform variant calling based on GATK best practices to identify SNPs and small indels. GATK-Variant Filtration was used to filter out variants with low calling quality (Phred QUAL<20) or low read depth (DP<20). Differences in SNP/indel frequency between the different strains were calculated for each SNP/indel, and those showing <50% difference were filtered out. Variants that were common to both vNF54 lines were crossed with transcriptomic data. Genome Browse (Golden Helix) was used to visualize alignments and variants, as well as for the detection of large subtelomeric deletions.

### ChIP-seq experiments and analysis

ChIP-seq experiments were performed essentially as previously described (53). In brief, chromatin was extracted from saponin-lysed synchronous parasite cultures at the late trophozoite / early schizont stage using the MAGnify Chromatin Immunoprecipitation System (Life Technologies) (34). After cross-linking and washing, chromatin was sonicated with an M220 sonicator (Covaris) at 10% duty factor, 200 cycles per burst, 140 W of peak incident power for 10 min. Next, 4 µg of chromatin were immunoprecipitated overnight at 4°C with 8 µg of antibody against H3K9me3 (Diagenode, C15410193) previously coupled to protein A/G magnetic beads provided in the kit. Samples were then washed, de-crosslinked and eluted following the MAGnify ChIP System recommendations but at all times avoiding high temperatures that could result in the denaturation of extremely AT-rich intergenic regions. De-crosslinking, proteinase K treatment, and elution were performed at 45°C (for 2 h, overnight, and 1.5 h, respectively).

Using a protocol adapted to a genome with an extremely high AT richness (56), libraries for Illumina sequencing were prepared from 5 ng of immunoprecipitated DNA. After end repair and addition of 3′ A-overhangs, NEBNext Multiplex Oligos for Illumina (NEB, E7335 and E7500) were ligated. Agencourt AMPure XP beads (Beckman Coulter) were used for the purification steps and the libraries were amplified (9 amplification cycles) with the KAPA HiFi PCR Kit (Kapa Biosystems) in KAPA HiFi Fidelity Buffer (5x). Finally, 0.9x AMPure XP beads were used to purify the amplified libraries and remove adapter dimers. After library size analysis using a 4200 TapeStation System (Agilent Technologies) and quantification using the KAPA Library Quantification Kit (Roche), sequencing was performed with a HiSeq2500 System (Illumina), obtaining 6 to 10 million 125 bp paired-end reads per sample.

Differential peak calling was performed using MACS2 (v. 2.2.7.1) following author’s recommendations. A first round of peak calling for every sample was performed using MACS2 “callpeak” command with parameters (-f BAMPE -g 2.41e7 --fe-cutoff 1.5 --nomodel --extsize 150) and then differential peaks were called using the “bdgdiff” command with parameters (-g 250 −l 300 --cutoff 5). Results where annotated using custom scripts in Python against the *P. falciparum* 3D7 annotation in PlasmoDB (v.46).

H3K9me3 coverage across the genome for each sample was calculated using the DeepTools (v.3.5.0) “BamCompare” command. After normalizing to RPKMs, coverage was defined as the log_2_(IP/input) and computed for 100 bp intervals. For each gene, we calculated the average coverage for the 1,000 bp upstream plus the first 500 bp of the coding sequence, because this is the region in which heterochromatin is typically associated with a silenced state.

### Determination of sexual conversion rates

Sexual conversion rates were measured by treating sorbitol synchronized ring-stage cultures with 50 mM GlcNAc (Sigma-Aldrich, A32869) for 5 days to eliminate asexual parasites. The sexual conversion rate was calculated as the gametocytaemia at day 5 relative to the initial rings parasitaemia at day 0, as previously described (57). Parasitaemia was measured by flow cytometry and gametocytaemia by light microscopy quantification of Giemsa-stained smears.

### Microsphiltration assay

In order to compare the deformability of erythrocytes infected with pNF54 and vNF54 (V63), we used a microsphiltration assay (58). Calibrated metal microspheres (96.50% tin, 3.00% silver, and 0.50% copper; Industrie des Poudres Sphériques) with two different size distributions (5-15 µm and 15-25 µm in diameter) were used to create a matrix within an inverted 1000 µl anti-aerosol pipette tip (Neptune). For this, 4 g of dry microspheres of each size range were resuspended in 12 ml of parasite culture medium (with 10% human serum) and 400 µl of this microsphere suspension was added into the tip and left to settle and form a 3-4 mm thick layer above the tip filter. Next, 600 µl of tightly synchronized 30-34 hpi cultures (1-6% parasitaemia) were placed on top of the microsphere layer, and then perfused through the microsphere matrix at a flow rate of 60 ml/h using an electric pump (Syramed SP6000, Arcomed Ag), followed by a wash with 5 ml of culture medium.

Samples collected after perfusion through the matrix (in triplicate) and the original culture (not passed through the matrix) were analyzed by flow cytometry. For this, 2 µl of erythrocytes pellet were collected in 200 µl of RPMI and washed twice with 200 µl of PBS, incubated for 25 min at room temperature with SYBR green (1:2,000 dilution of Sigma S9430 solution), washed twice again with 200 µl of PBS and finally resuspended in 1.5 ml of PBS. The parasitaemia of each sample was then measured with a BD Accuri C6 cytometer, with 100,000 events recorded per sample.

### Sorbitol assay

To test sorbitol sensitivity, tightly synchronized 20-24 hpi pNF54 and vNF54 (V63) cultures were treated with a sorbitol-containing isosmotic solution (300 mM sorbitol supplemented with 10 mM HEPES, 5 mM glucose and adjusted to pH 7.4) or the same isosmotic solution with 100 µM of the general anion channel inhibitor 5-Nitro-2-(3-phenylpropylamino) benzoic acid (NPPB, Sigma-Aldrich), which inhibits malarial new permeation pathways (59). Parasitaemia was determined before (time 0) and after (time 60’) incubation at 37°C for 1h, using flow cytometry. The lysis percentage was calculated for each condition using the following formula: %lysis = [1-(parasitaemia time 60’ / parasitaemia time 0’)]*100.

### Western Blot

Synchronous cultures containing mainly ∼40 hpi schizonts were purified by magnetic isolation, divided into pellets of approximately 2.5×10^6^ schizonts and stored frozen at −80°C. After thawing, proteins were denaturized in SDS-PAGE protein loading buffer for 5 minutes at 95°C and resolved by SDS-PAGE on 4-12% bis-tris Criterion XT Precast gels (Bio-rad), transferred to a PVDF membrane and blocked for at least 1h in 1% Casein blocking buffer (Sigma). Membranes were incubated overnight at 4°C with the following primary antibodies: purified mouse antiserum against a PfMC-2TM protein (PF3D7_0114100) at 1:200 (kindly provided by Catherine Braun-Breton) (60) and mouse anti-HSP70 antibody (61) at 1:2,000. Samples were then incubated for 1h at room temperature with horseradish peroxidase-conjugated anti-mouse IgG secondary antibody (Promega) at 1:10,000 and peroxidase was detected using the Pierce chemiluminescence system (Pierce) following the manufacturer’s instructions. To control for equal loading, parts of the membranes corresponding to different molecular weight ranges were separately hybridized with different antibodies. Signal quantification was performed using ImageJ software.

## Data availability

The microarray and ChIP-seq data have been deposited in the GEO database with accession numbers GSE166258 (token for reviewer access: kjulwgmorrqvxof) and GSE166390 (token for reviewer access: ihczaeamxxmffmh), respectively. ChIP-seq data can be visualized at the UCSC genome browser using the following link: http://genome.ucsc.edu/s/apickford/apickford_lmichel. Whole genome sequence data has been deposited at the Sequence Read Archive (SRA) database with accession number PRJNA699845 (reviewers can access the data using the following link: https://dataview.ncbi.nlm.nih.gov/object/PRJNA699845?reviewer=25eaj128nc3s4bjdjg9ji4v4f1). The scripts used for the analysis of microarray and ChIP-seq data will be made available upon reasonable request, without any restrictions.

## Acknowledgments

We thank Catherine Braun-Breton (Université de Montpellier) for providing antibodies against a PfMC-2TM protein; the Genomics Unit at the CRG for assistance with genome sequencing; Gloria P. Gómez-Pérez, José Muñoz (ISGlobal) and the volunteers and clinical staff who participated in the CHMI in Barcelona; Sofía Mira-Martínez and Ariel Magallón-Tejada (ISGlobal) for their contribution to the collection of samples for transcriptional analysis in the CHMI study; and Stephen L. Hoffman and Kim Lee Sim (Sanaria) for providing cryopreserved *P. falciparum* sporozoites. This work was supported by grants from the Spanish Ministerio de Ciencia e Innovación (MCI)/ Agencia Estatal de Investigación (AEI) [SAF2016-76190-R and PID2019-107232RB-I00 to A.C.], co-funded by the European Regional Development Fund (ERDF, European Union), and grants by the Instituto de Salud Carlos III and Fundación Ramón Areces to P.L.A. for the CHMI trial. F.D. and C.L. were supported by CNRS, Inserm, and the Fondation pour la Recherche Médicale under award number “Equipe FRM” EQ20170336722D. A.K.P. is supported by a fellowship from the Secretary for Universities and Research, Catalan Government (FI_B 00373), co-funded by the European Social Fund (ESF), European Commission. L.M.-T. is supported by a fellowship from the Spanish Ministry of Economy and Competitiveness (BES-2017-081079), co-funded by the European Social Fund (ESF). Our research is part of ISGlobal’s Program on the Molecular Mechanisms of Malaria, which is partially supported by the Fundación Ramón Areces. We acknowledge support from the Spanish Ministry of Science and Innovation through the “Centro de Excelencia Severo Ochoa 2019-2023” Program (CEX2018-000806-S), and support from the Generalitat de Catalunya through the CERCA Program.

